# The Induced Fatigue on Attention Networks of Active and Inactive Individuals Effect of Exercise

**DOI:** 10.1101/2023.08.14.553215

**Authors:** Maryam Kayvani, Akram Kavyani, Sana Soltani

## Abstract

**Background:** The relationship between physical activity and cognitive functions is well established, with regular exercise improving cognitive functions. However, there is less clarity surrounding the effect of exercise-induced fatigue on attention network particularly in those with a history of continuous physical activity.

**Aims:** This research was aimed to determine whether exercise-induced fatigue can affect cognitive functions, especially those involved in attentional control (i.e., alerting, orientation, and executive functions) and to identify any differences in attentional control between active and inactive individuals after exercise-induced fatigue.

**Methods:** We compared the performance of 24 physically active and inactive participants in the Attentional Network Task, which allows for the assessment of the executive, orienting and alerting networks. Under two conditions regarding exercise-induced fatigue (pre-fatigue and post-fatigue), we used sub-maximum aerobic endurance training to induce fatigue to the exhaustion point.

**Results:** The results showed that fatiguing exercise improved alertness in both groups; however, the executive control network of the active group improved while the orienting and executive control networks of the inactive group performed worse.

**Conclusions:** Depending on the participants’ degree of physical activity and the particular task used to test each of these attention networks, exercise-induced exhaustion had a different impact on different attention networks.

## Introduction

Muscular fatigue is an exercise-induced reduction in muscle ability (1), It originates at two levels of the motor pathway: peripheral and central (2). Peripheral fatigue motion is caused by changes to the muscle itself, typically at the neuromuscular junction. Central fatigue is caused by changes in the central nervous system (CNS), involving the brain. The level of neural signalling to the muscles decreases, consequently impairing the ability of the muscle to perform its normal functions. (1, 3)The effect of motion-induced fatigue on cognition and the interrelation between CNS (central nervous system) circuits of physical fatigue and cognitive functions have rarely been studied. Studies on the effect of mental fatigue have shown that it has a detrimental effect on multiple factors, such as behaviours, physiology, and psychological outcomes during exercise. Notably, a significant contributor to the harmful effect of mental fatigue on endurance performance has been higher perceived exertion(4). There has been no association estabilished between mental and central fatigue(5). Despite studies on the effect of mental fatigue on physical function, the reciprocal relationship of physical function effect on mental fatigue remains unclear.

Descriptive reviews (6, 7) and meta-analytic studies (8-11) have reported conflicting results regarding exercise effects on cognitive function. Some studies have focused narrowly on the short-term effects of exercising on cognitive function, as, for example, a study that assessed the cognitive performance of 15 cyclists immediately after one training session and showed improved simple reaction time and executive function but no change in selective reaction time or finger tapping speeds (12). However, Grego et al. (2004) found a speed reduction in stimulus classification and information processing among young, actively trained, male cyclists. Similarly, other investigators examined the effect of a 60-min cycling exercise on cognitive functions and found a subsequent decrease in complex perceptual discrimination and an increased reaction time on the memory-based vigilance test, while detection and identification did not differ between exercise and control groups (13). Wilke et al. (2019) concluded that the acute effect of resistance exercise (RE) on cognitive function depends on cognitive sub-domains. Considering the specific cognitive domains or task requirements, working memory and attention remained unaffected whereas inhibitory control and cognitive flexibility was improved after RE(11). It is worth noting that most studies explore the effects of exercise on cognitive function rather than the effects of fatigue after exercise. The effects of exercise-induced fatigue on cognitive functions may also be task-specific. In addition to different exercise effects on varied cognitive tasks, the effect of physical induced fatigue should be considered. Furthermore, evidence for the effects of intense exercise on cognitive control has mostly come from studies using incremental protocols in which the effects of exercise are confound with the effect of exhaustion(14, 15). The present study aimed to separate the effects of exercise-induced fatigue from the exercise intensity using a maximal voluntary contraction (MVC) test immediately after exercise and measure cognitive function thereafter.

Given the importance of task type in research on exercise-induced fatigue and the basic role of attention for most cognitive and motor tasks, it is essential to evaluate the effects of attentional tasks, particularly . The term executive functions (EFs) refers to a wide range of top-down mental processes needed for the cognitive control of behaviors. These wide ranging cognitive skills have often been reduced in exercise research to three core EFs: inhibition (inhibitory control and interference control), working memory, and cognitive flexibility (16). Based on these core EFs, other higher-order EFs such as reasoning, problem solving and planning are built (17, 18). Inhibitory control of attention (a core EF) enables us to selectively attend by suppressing attention to particular stimuli, based on our intention (19). Selective attention initiates many cognitive processes, includes alerting, and orienting, and involves executive control neuronal networks in the brain (i.e., Attention Networks or Ans); (20). Alerting is either tonic (over sustained periods) or phasic (temporarily increased readiness in response to a warning) (20, 21). The orientation network changes either attention to specific spatial situations or prioritizing of some sensory inputs for processing (20, 22). The executive control network identifies incompatibilities and inhibits annoying information in top-down processes (23, 24). The Attention Network Test (ANT) measures these attentional networks’ (AN) functions (25, 26). Through an executive control, which is one of AN function with an essential role in sports, one plans, makes decisions, finds errors, gives new responses, or overcomes habitual actions. executive function can also be generalized to other cognitive tasks (27). However, the executive control of ANT can be named conflict inhibition, and it is assessed by subtracting the mean reaction time from trials with congruent flankers from that of trials with incongruent flankers (26). Although the three attentional networks appear functionally and anatomically independent, t here is functional integration and interaction among these three attentional networks. The alerting network seems to inhibit the executive function network, the orienting network influenced the executive function network in a positive way and alertness increased orienting(28, 29). Therefore, considering these interactions, the use of ANT in exercise-induced fatigue seems more logical than separate executive function tests.

In order to understand inconsistencies in the results of past research regarding relationships between exercise-induced fatigue and cognitive functioning, we must consider the influences in past studies of failures to measure fatigue in fatigue-inducing protocols, different cognitive tasks and cognitive task scheduling, and even the participants’ varied physical fitness. Variations in physical fitness as a results of participation in open and close sports have had different effects on cognitive function (30), and especially executive functions (31, 32), and may cause varied physiological changes in fatigue mechanisms and perception. The exact extent of fatigue in active and inactive participant research groups has rarely been studied.

It is essential to study this relationship because fatigue levels can be found in inactive individuals, whose lower physical condition makes them more susceptible to fatigue. There is a well-established relationship between physical activity and cognitive functions where regular physical activity improves cognitive functions. (6, 33).

In present study, we investigated the effects of exercise-induced fatigue on the AN functioning of active and inactive young individuals. For this purpose, the fatigue-inducing protocol for both groups were designed for participants individually, according to their beginning physical fitness levels. To determine if our experimental manipulation induced fatigue, we measured MVCs pre-post intervention. After inducing performance decline in both groups, we examined the effect of physical exercise-induced fatigue on their AN function, as measured by ANT.

## Methods

### Participants

#### Ethic Statement

Twenty-four young (19 females, 5 males), healthy adults signed informed consent forms to participate in this study following a full explanation of the research protocol. We received advanced approval to conduct this research after its review by the Biological Research Ethics Committee of SB University, and receipt of the Ethics confirmation code IR.SBU.REC.1398.123 that was issued for the research. The recruitment period for this study commenced on 22nd June 2019 and concluded on 23rd August 2019, as specified.

#### Participants

Twenty-four young (19 females, 5 males), were equally divided into active and inactive groups based on their responses to the international Physical Activity Questionnaire-Long Form (IPAQ) (34).

Based on IPAQ, there are three suggested levels of physical activity for population classific ation: 1) Low/ Inactive, 2) Minimally Active, 3) HEPA active (health enhancing physical activity; a high active category). The criteria for these three levels are presented below.

Inactive (CATEGORY 1)

This is the lowest level of physical activity. Individuals who do not meet the criteria for category 2 or 3 are considered ‘insufficiently active’.

2. Minimally Active (CATEGORY 2)

The minimum activity pattern that can be classified as ‘sufficiently active’ is any one of the following 3 criteria:

a. 3 or more days of vigorous activity of at least 20 minutes per day OR
b. 5 or more days of moderate-intensity activity or walking of at least 30 minutes per day OR
c. 5 or more days of any combination of walking, moderate-intensity or vigorous intensity activities achieving a minimum of at least 600 MET-min/week.

HEPA active (CATEGORY 3)

The two criteria for classification as ‘HEPA active’ are:

a. vigorous-intensity activity on at least 3 days achieving a minimum of at least 1500 MET-minutes/week OR
b. 7 or more days of any combination of walking, moderate-intensity or vigorous intensity activities achieving a minimum of at least 3000 MET-minutes/week

Those who scored category 2(n=3) and 3(n=8) were considered active. Participants of Active groups played at different sports such as football(n=3), futsal(n=1), water polo(n=1), volleyball(n=3), and basketball(n=3). Individuals who scored category 1 were placed in the inactive group. Both groups had matching ages. All participants were right-handed and had a normal or corrected-to-normal vision.

Lower extremity muscles, particularly knees, ankles, and supporting muscles, were examined to ensure participants had no previous injuries, surgery, congenital abnormalities, or stroke. Individuals were asked to maintain their eating and resting routines for at least 2 days before the sessions, drink sufficient fluids, and avoid alcoholic and caffeinated beverages, psychotropic drugs, and antidepressants 24 hours before sessions.

#### Instruments

Instruments included the Iinternational Physical Activity Questionnaire-Long Form (IPAQ)(34), Mini Mental State Exam (MMSE) Cognitive Health Questionnaire (35), isokinetic dynamometer to measure maximal muscle strength(Biodex Medical Systems, Inc., Shirley, NY, USA), Monark ergometer bike designed by Monark Exercise AB, Sweden, Borg rating of perceived exertion, Attention Network Test (ANT) (29), measuring tape, a scale to measure anthropometric characteristics, Snellen chart to measure visual acuity, and a “14” laptop for presenting tasks.

#### Procedure

This research was conducted in two 90-minute sessions. During the first session, the participants took the MMSE followed by the ANT for 20 minutes under normal physical conditions and without fatigue. Maximal contractions in the knee flexor (hamstring) and extensor (quadriceps) muscles were also measured at a 90-degree angle to familiarize participants with the protocol for measuring maximal contraction.

The second session took place within 1 to 3 days after the first session. Participants were asked to avoid tiring physical or cognitive exercise and to maintain their daily routine between the two sessions.

At the beginning of the second session, all participants completed the MVC test. After that a stationary bike was used for the exercise-induced fatigue protocol. Next, fatigue was measured by maximal voluntary contraction (MVC) using an isokinetic dynamometer. Both groups then performed the ANT and MVC tests to measure fatigue from reduced muscle strength according to the fatigue protocol. An overall view of the protocol can be found Fig 1.

**Fig 1.**
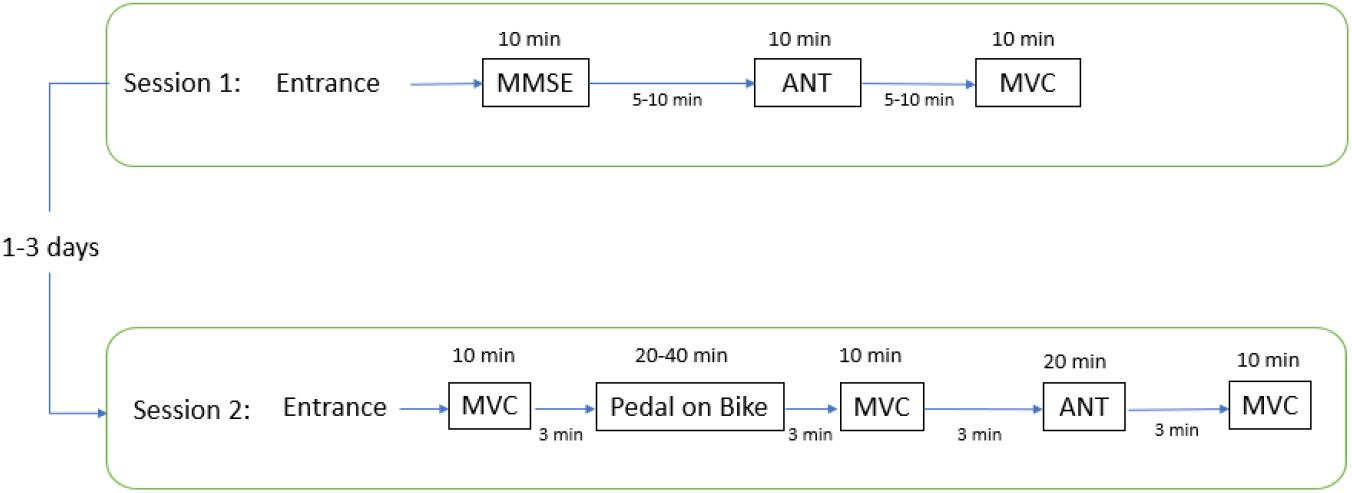
Overview of experimental protocol.

#### Maximum Voluntary Contraction (MVC)

Knee flexors (hamstrings) and extensors (quadriceps) MVC was measured using an isokinetic device during an isometric contraction before and after the fatigue protocol.

During the test, participants sat in an adjustable chair with their hips and knees bent at 90 degrees, their ankles in a neutral position, their pelvis and chest taped in place, and a monitor placed 50 cm away from them was, and a visual representation showed feedback about the degree of contraction. Participants were instructed to generate three maximal voluntary contractions of knee flexors and extensors (5 seconds each) with 2 minutes rest between each. They were encouraged to exert maximal effort and contraction on all three trials. In the first session, they were asked to perform three knee flexors and extensors contractions to learn how to properly perform the MVC test.

#### Fatiguing Exercise

We aimed to design an endurance exercise protocol to assure that participants stopped exercising at exhaustion. Since we planned to compare active and inactive participants, progressive protocols were used to ensure equal fatigue and homogeneous exercise abilities in terms of VO2Max.

Participants performed dynamic physical exercise using a Monark ergometer to induce global fatigue. They first adjusted the saddle and handlebars to their preferred position and then pedaled at their preferred speed for 10 minutes to warmup and familiarize themselves with the task. They continued according to ramp exercise protocol until exhaustion. Voluntary exhaustion ensured that the practice load was personally optimized for each participant, and the total time of exercise varied based each personal physical fitness level. The initial load was 25W, adding 5W every 5 minutes. The total time of exercise was varied based each personal physical fitness level. The visual feedback displayed allowed the participants to maintain the desired pedaling speed of 50-60 rpm. They were encouraged to maintain maximum engine power. Each exercise would end at the participant’s discretion or when their pedaling speed was no longer stabilized at speeds over 50 rpm. Voluntary exhaustion was defined as the point where participants voluntarily stopped or when they could no longer maintain a pedaling frequency above 50 rotations per minute (RPM) more than 10 s despite strong verbal encouragement and they accrued ratings of perceived exertion of 19-20.

#### Attention Network Test Protocol

ANT was measured using a standard 20-minute version of the test designed by Fan et al. in 2009 (29). ANT was presented and the data were recorded by Psych toolbox. The details of ANT are shown in Fig 2. In this task, each trial begins with a central fixation point followed by one of three cue conditions (no-cue, double-cue, spatial-cue). A target display appears 400 ms after the offset of the cue. The target condition is congruent or incongruent. Details of cue and target display times and cue target intervals are shown in Fig 2. To measure participants’ attentional function, i.e., alerting, orienting, and executive control, randomly equal attempts made in all blocks. The test instructions emphasize maximum speed and accuracy. Participants completed a practice block with 24 full feedback trials. The test phase consisted of 288 trials, split into three blocks of 96. Finally, the reaction time and performance accuracy data were automatically recorded.

**Fig 2.**
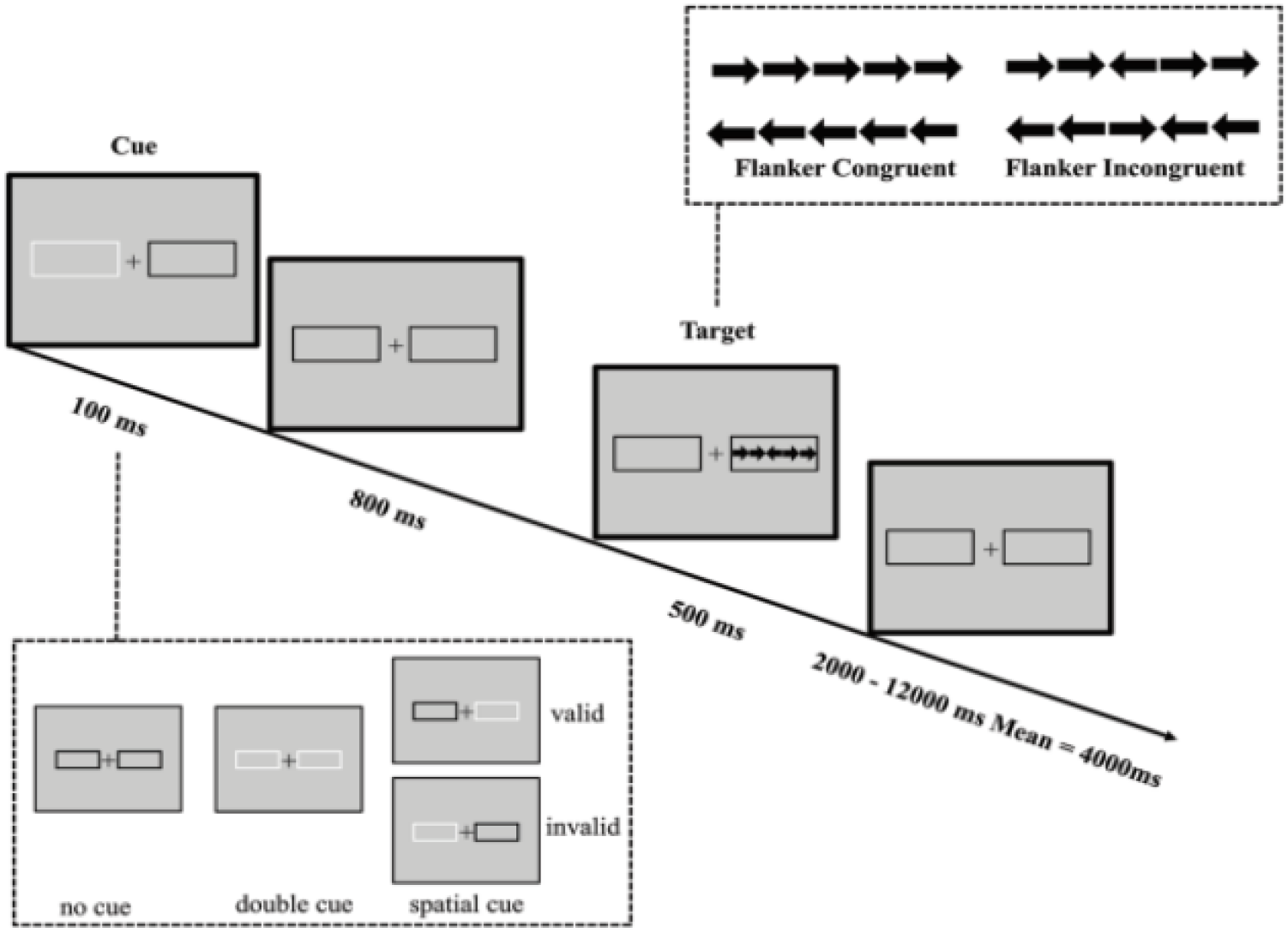
The Attention network Test (ANT) experimental procedure.

**Fig 3.**
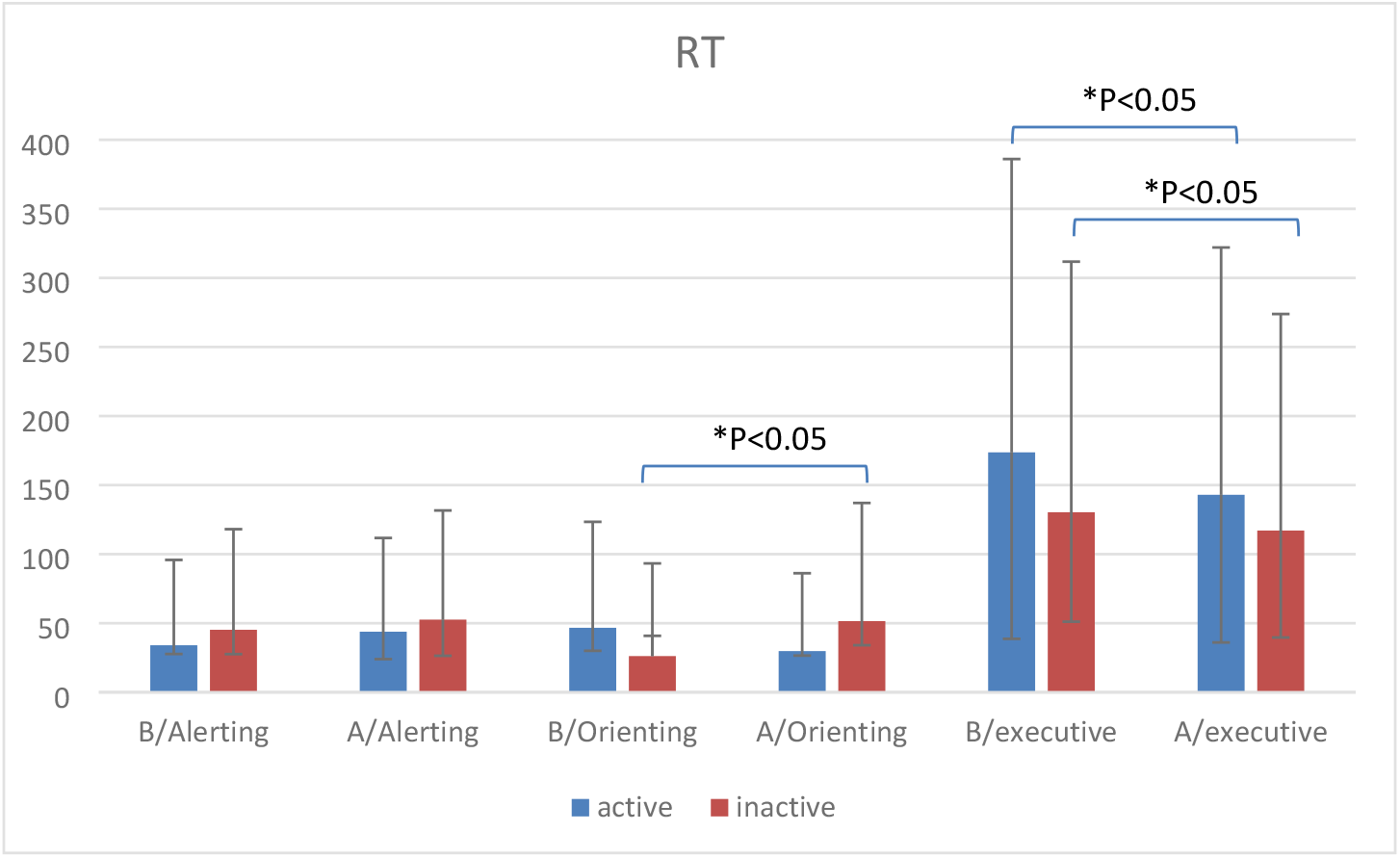
Before(B) and After(A) fatigue scores of Attention Network index in active and inactive participants for Reaction Time (RT). Note: * P<0. 05.

**Fig 4.**
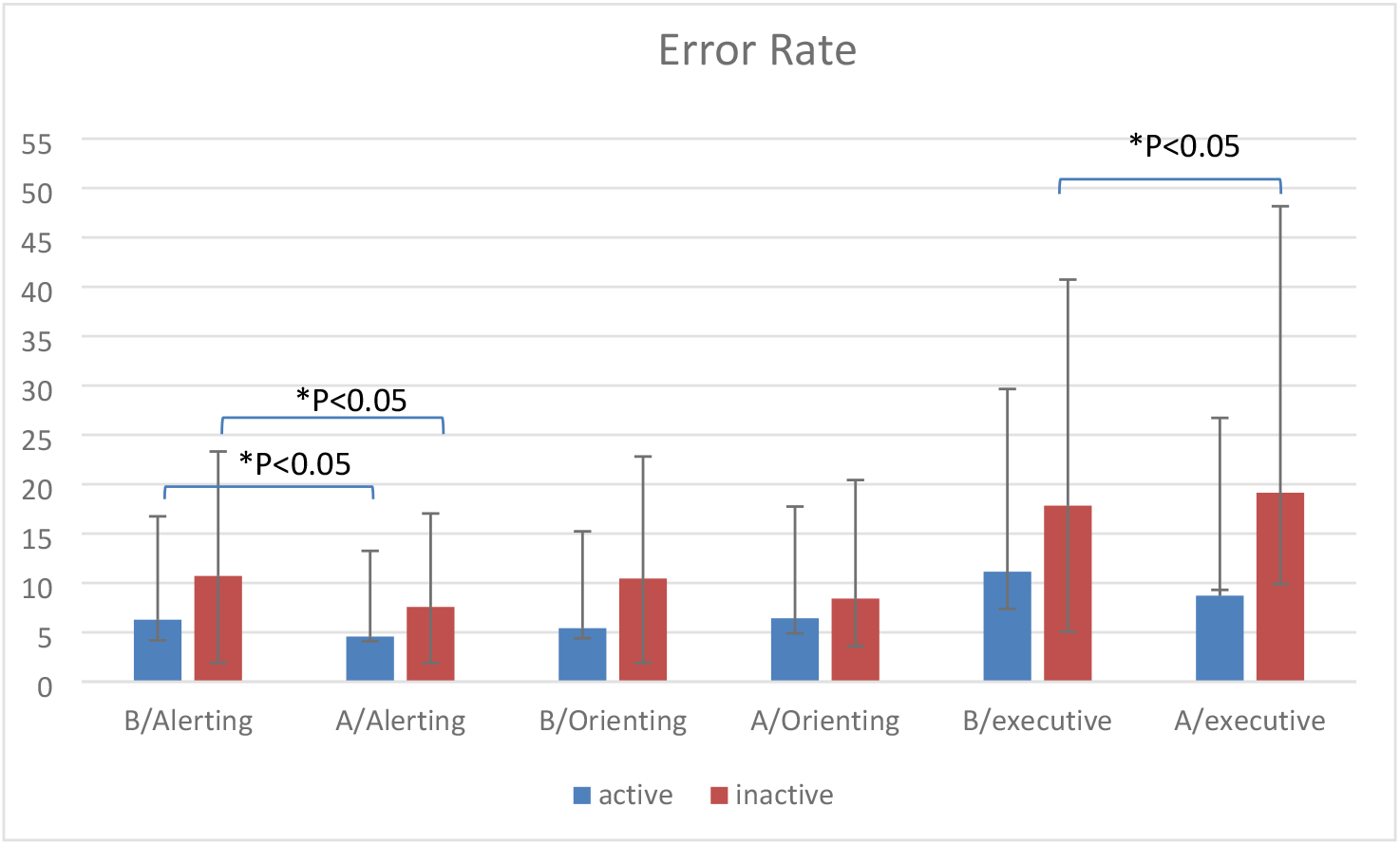
Before(B) and After(A) fatigue scores of Attention Network index in active and inactive participants for Error Rate (RT). Note: * P<0. 05.

### Statistical analysis

Descriptive statistics of frequency, mean, standard deviation descriptive statistics were used to describe the data. The Shapiro-Wilk-Test confirmed that the data was normally distributed and there was homogeneity of variance (Levin’s test). We then we tested the hypothesis using a two-way (group * condition) (2 * 2) analysis of variance (ANOVA).

A priori power analysis was performed to estimate sample, based on data from published study Moore et al.,(13) and predicted effect size of 0.14 for changes in a simple and complex perceptual task after fatigue. We consider the least effect size in their study that was significant. With an alpha = 0.05 and power = 0.95, the predicted sample size needed with this effect size (GPower 3.1 software) was approximately N = 24 for a within-between ANOVA test (two measurements, two groups). Therefore, our proposed sample size of 24 was appropriate for the main objective of this study. All outcome measures demonstrated adequate internal consistency, with inter-rater reliability coefficients ranging from 0.65 to 0.87 for all measures. The software employed was SPSS Statistics 20.0,, and G.Power 3.1.

## Results

Table 1 shows the descriptive data on the age, height, weight, MMSE score, IPAQ score, Borg rating score, and pre-fatigue to post-fatigue MVC ratio of the groups. According to the table, the reduction in maximal contraction force of both groups was 25%, indicating the effectiveness of the fatigue protocol.

**Table 1.**
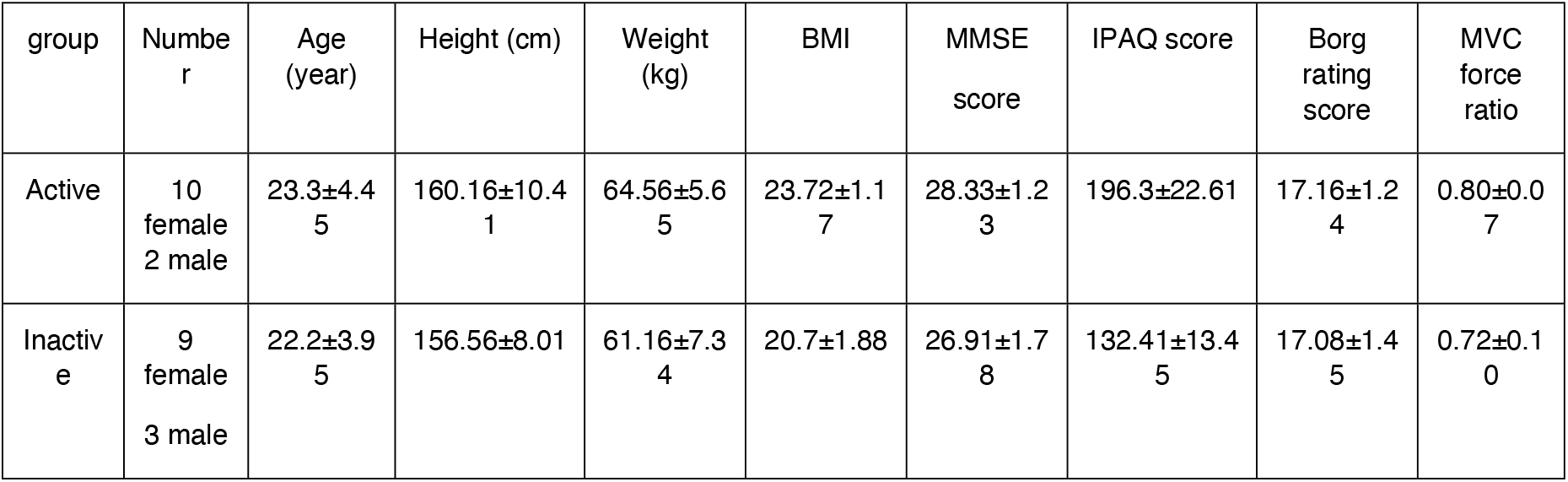
Descriptive Data on Age, Height, Weight, MMSE score, IPAQ score, Borg rating score and Pre-Fatigue to Post-Fatigue MVC Ratio of Groups.

Table 2 illustrates the mean and standard deviation of ANT reaction time and error of active and inactive groups in pre-and post-fatigue conditions. The efficiency of alerting was calculated using RTs with no cue minus RTs with double cues, orienting was RTs with central cues minus RTs with spatial cues, and executive function was RTs of incongruent flankers minus RTs of congruent flankers.

**Table 2.**
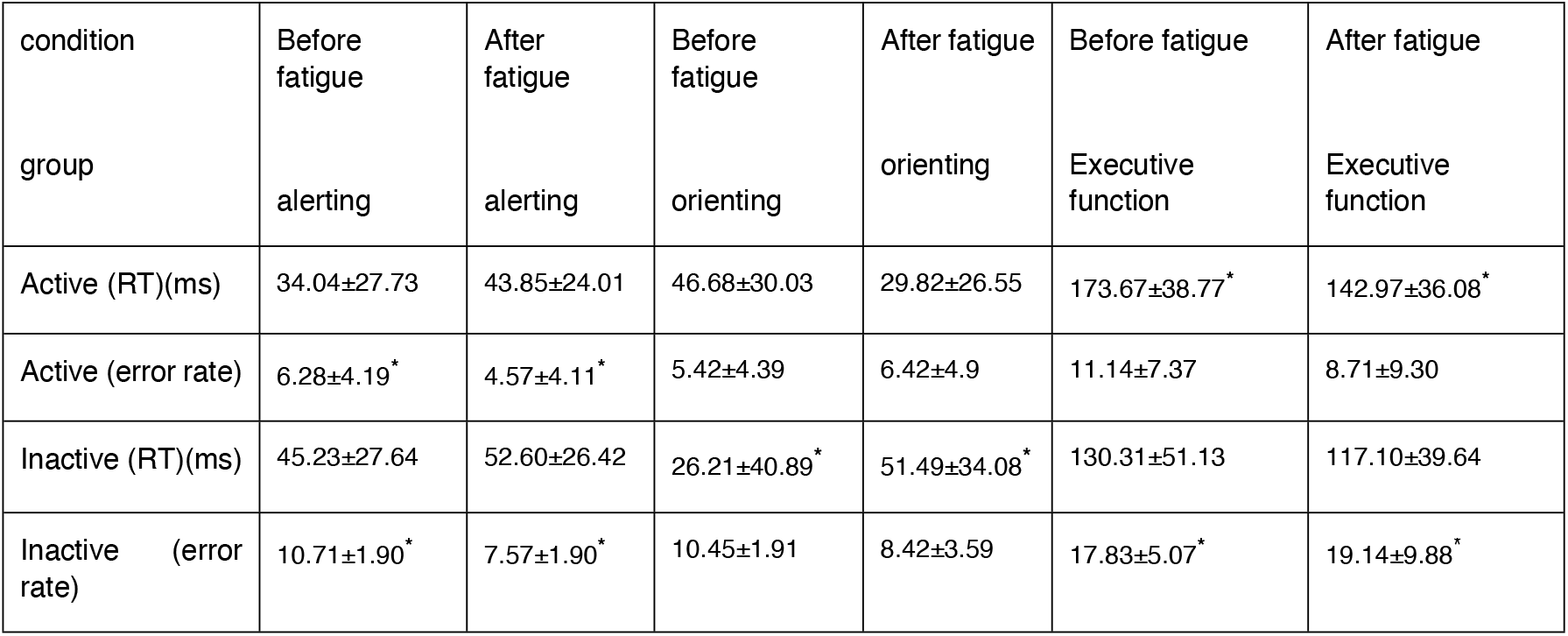
Mean and Standard Deviation of Attention Networks Reaction Time and Error of Pre- and Post-Fatigue Conditions in Active and Inactive Groups. * Indicates a Significance Difference (p< 0.05) between Pre- and Post-Fatigue Conditions.

| condition | Before fatigue | After fatigue | Before fatigue | After fatigue | Before fatigue | After fatigue |
| --- | --- | --- | --- | --- | --- | --- |
| group | alerting | alerting | orienting | orienting | Executive function | Executive function |
| Active (RT)(ms) | 34.04±27.73 | 43.85±24.01 | 46.68±30.03 | 29.82±26.55 | 173.67±38.77* | 142.97±36.08* |
| Active (error rate) | 6.28±4.19* | 4.57±4.11* | 5.42±4.39 | 6.42±4.9 | 11.14±7.37 | 8.71±9.30 |
| Inactive (RT)(ms) | 45.23±27.64 | 52.60±26.42 | 26.21±40.89* | 51.49±34.08* | 130.31±51.13 | 117.10±39.64 |
| Inactive (error rate) | 10.71±1.90* | 7.57±1.90* | 10.45±1.91 | 8.42±3.59 | 17.83±5.07* | 19.14±9.88* |

In the next step, a 2 ×2 ANOVA was used to analyze <u>reaction time</u> and <u>error scores</u>, consisting of group (active and inactive) and the condition (before and after fatigue) factors. According to the alerting network data, the main effect of group (F (1, 22) = 1.59, p = 0.22, η^2^ = 0.06), the main effect of the condition (F (1, 22) = 1.36, p = 0.26, η^2^ = 0.05), and the interaction effect of the group × condition was not significant (F (1, 22) = 0.03, p = 0.86, η^2^ = 0.001) for <u>reaction time</u> variable. The <u>error variable</u> in the alerting network showed a significant main effect (F (1, 22) =5.38, p=0.03, η^2^ = 0.31). In other words, the error scores were significantly higher in the inactive group than in the active group. The main effect of condition was also significant, (F (1, 22) =10.50, p=0.007, η^2^ = 0.46) and the error score was higher after fatigue. Interaction effect of condition × group was not significant in this case (F (1, 22) =0.90, p=0.25, η^2^ = 0.07). P<0.05

According to orienting network data, the main effect of group (F (1, 22) = 0.15, p = 0.69, η^2^ = 0.007), and the main effect of condition (F (1, 22) = 1.40, p = 0.24, η^2^ = 0.06) were not significant, but the interaction effect of the group × condition was significant (F (1, 22) = 4.27, p = 0.05, η^2^ = 0.16) regarding <u>reaction time variable</u>. <u>Reaction time</u> after fatigue increased significantly compared to the pre-fatigue conditions of the inactive group according to the paired t-test (t = 2.37, df = 11, p = 0.03, MD = 25.28). <u>Error variable</u> in orienting network (Table 2) showed a significant main effect (F (1, 22) =4.67, p=0.05, η^2^ = 0.01). In other words, <u>error scores</u> were significantly higher in the inactive group compared to the active group. However, interaction effect of condition × group was not significant in this case (F (1, 22) =0.06, p=0.79, η^2^ = 0.006).

According to the data analysis of executive function network, the main effect of group F (1, 22) = 4.74, p = 0.04, η^2^ = 0.18) and the main effect of condition (F (1, 22) = 8.92, p = 0.007, η^2^ = 0.28) were significant but the interaction effect of the group × condition was not significant (F (1, 22) = 1.52, p = 0.23, η^2^ = 0.06) for <u>reaction time variable</u>. Generally, after fatigue, both groups had lower reaction times in the executive task. <u>Error variable</u> in executive function network showed a significant main effect (F (1, 22) =6.19, p=0.02, η^2^ = 0.34). In other words, <u>error scores</u> of the inactive group were significantly higher than the active group.

Condition main effect (F (1, 22) =0.28, p=0.60, η^2^ = 0.02) and interaction effect of condition × group (F (1, 22) =0.22, p=0.64, η^2^ = 0.01) were not significant. The inactive group responded faster but less accurately in the fatigued condition.

## Discussion

This study aimed to determine whether exercise-induced central fatigue affecting the central nervous system can alter cognitive functions, particularly alertness, orientation, and executive functions involved in selective attention of active and inactive individuals.

The active and inactive groups did not show different reaction times for alerting under pre-and post-fatigue conditions. Inactive individuals had more errors (lower accuracy) compared to active individuals in the pre- and post-fatigue conditions, and their post-fatigue alerting errors were reduced, i.e., they were more accurate in post-fatigue. Regarding orientation, the reaction time of the inactive group in the fatigue condition was significantly increased. Regarding the executive functions network, both groups responded faster in the fatigue condition, but the inactive group had more errors, indicating lower accuracy than the active group. Thus, physical activity with subsequent fatigue improved alertness in both groups decreased the orientation speed of the inactive group, and had no effect on the orientation speed in the active group. The executive control network improved in active individuals, but the executive control network error was increased in inactive individuals after fatigue.

Our results replicated the general facilitating effect of exercise on alerting network by an improvement in the general state of tonic vigilance and preparedness. Interestingly, no significant differences were observed between the active and inactive groups with regard to the alerting network. These results may be clarified somewhat by the theory of arousal and exercise-induced increases in adrenaline-noradrenaline secretion. These results are partially in line with the finding of recent meta-analysis(36), which shows that the performance of tasks involving quick decisions and automated behaviors improves with exercise-induced arousal.

These results were consistent with those of Huertas et al., and Hogervorst et al., who examined the effects of an exercise session with different intensities on the cognitive performance of professional cyclists and concluded that short-term exercising reduced the alerting effect but other AN functions were **unaffected**, so orientation and executive function. According to the authors, short-term exercise may have altered the phasic awareness by increasing the level of tonic vigilance. In the present study, the alerting accuracy score improved after physical activity. Unlike Huertas et al., fatiguing physical activities disrupted the orientation network in inactive individuals of the present study and improved executive functions, especially in active individuals. This difference is likely due to different levels of fitness and intensity of physical activity(37, 38).

In regard to the effect of exercise-induced fatigue on the orienting network, our results showed that the level of physical activity modulated this function inducing a decrease in visuospatial attention in inactive individuals. We argue that normal exogenous attentional function was affected by exercise-induced fatigue, which affected inactive participants more than active participants. As a consequence, target processing was prioritized over irrelevant stimuli. Similar to the study conducted by Moore et al., the effect of orientation decreased in the present study. These researchers also showed a decrease in complex perceptual differentiation tasks after 60 minutes of cycling. Unlike the present study, they also found an increase in the reaction time during the memory-dependent vigilance test (13). Therefore, the effect of exercise-induced fatigue may be task-specific and have a more significant impact on the perception task, which requires relatively automated processing compared to the effort-based memory task. However, and in line with previous studies (37, 39) the present results did not show changes in the orienting speed of active individuals after exercise or subsequent fatigue.

More importantly for the purpose of the current study, our results similar to others (40), revealed that the effect of exercised-induced fatigue on executive function was moderated by the level of physical activity. Adaptions to physical training improved executive control in active participants when they were fatigued, but conversely, lack of physical activity induced a weaker executive function in inactive participants in the fatigued state. The executive function is necessary to detect the final target related to the limited capacity of the attentional system. Higher levels of physical activity could result in more efficient focal attention processing and top-down processing, as well as better self-regulation and inhibitory control in active individuals. These results are inconsistent with findings of Hilman, Kramer, Belopolsky, and Smith as well as Boucard et al., who showed that long-term regular exercise does not affect inhibitory functions (as a subset of executive functions)(41, 42). This difference is likely due to different definitions of active and inactive groups, as well as different measurement indexes of executive functions in various tasks. Unlike the present study, previous researches have probably used only the reaction time index regardless of accuracy. Furthermore, the complicated and multidimensional nature of executive functions and the difficulty in making the measurement of executive functions operational could also contribute to the inconsistencies in the results. The results of the present study also did not agree with the results of Tomporowski et al., who found a decline in executive functions after exercising in dehydrated conditions (43). The difference in results can be mainly attributed to the participants’ readiness levels, the type of activity that induces fatigue and, more importantly, the dehydrating conditions.

More interestingly, even though the independence of attention network functions has been established Xuan et al., factors out that there may be an interplay amongst attention networks, which means that they’re moreover interrelated(44). In line with this idea, our results indicated the interaction effect of alerting by executive control. But obviously, the degree of cognitive adaptation of individuals to physical activity affects this relationship. In other words, the increased sensitivity to exogenous stimuli due to increased arousal after physical activity, which was directly characterized by improved post-fatigue alertness in both groups, is related to the lack of ability to select relevant information in inactive individuals and is therefore reduced executive function of the inactive individual. Rather, improved alertness result in better orientation and prioritization of relevant information in the orienting network, making target detection in the active individuals more efficient. Thus, the active individuals had a better executive function after fatigue. It is possible that the adaptation created by the regular activity has led to the arousal level of the active group within the desired arousal level for those individuals. Similar to this study, Hogervorst E, Riedel W, Jeukendrup A, Jolles J. found that alertness and executive functions of the trained cyclists were improved after training (38).

This study had unique characteristics setting it apart from other similar studies. First, we examined the effects of endurance exercise-induced fatigue on cognitive functions, especially on AN. Previous researches had investigated the effect of short-term or long-term exercise regardless of the effect of physical fatigue on AN. Second, the MCV test was used for measuring post-exercise fatigue objectively. However, physical fatigue degree had not been objectively measured in studies that examined the effects of fatigue on cognitive functions. In this study, the MVC was reduced by nearly 25% after physical activity. Finally, this study examined how regular exercise adaptation impacted the effect of exercise-induced fatigue on the cognitive functions of active and inactive groups. In previous studies(39, 40, 45), the effect of exercise-induced fatigue on cognitive functions was examined only in active and professional individuals. It is important because exercise has been proved to stimulate the brain to maintain functions and structures involved in executive functions(2, 46, 47). This feature is achieved through relatively long-lasting physiological changes made in the structures of executive control.

Interpretation of the results was limited due to the difficulty in separating the beneficial effects of short-term aerobic exercise on the cognitive system from the adverse effects of aerobic exercise-induced central fatigue. A further study comparing the effects of exercise-induced fatigue and the effects of non-fatiguing aerobic exercise is needed to clarify any ambiguity in the interpretation of these results. When researchers are able to measure cognitive functions, mainly executive functions, during physical activity, the effect of physical fatigue on cognitive functions becomes more visible.

Other non-physiological factors may also be involved in exercise-induced fatigue in executive function. For example, open-ended exercises that require increased cognitive processing have been found to show selective benefits for improving inhibitory control(30-32). It is possible that the athletes included in the present study who participated in open sports exhibited a better level of executive function than the passive participants and that this may have had an impact on tasks that require a relatively high level of executive function. Therefore, an important direction for future work would be to study the effects of exercise-induced fatigue on cognition, taking into account different sports experiences (e.g., open skilled sport versus close skilled sport).

## Acknowledgments

The authors would like to thank all participants in this study.

## Availability of data and material

All data were presented in the main manuscript or additional supporting files. The raw data used and/or analyzed during the current study are available upon reasonable request to the corresponding author (MK).

